# A second-generation “hypoxia in a pill” rescues neurodegenerative phenotypes across distinct mouse models

**DOI:** 10.64898/2026.06.30.734089

**Authors:** Hong Wang, Eizo Marutani, Luca Zazzeron, Marissa Menard, Laura Volpicelli-Daley, Fumito Ichinose, Vamsi K. Mootha

**Author notes:** Corresponding author: Vamsi K. Mootha, M.D.

## Abstract

A growing body of pre-clinical research is demonstrating the therapeutic potential of chronic, continuous hypoxia (11% FIO2) for rare and common diseases (1). However, the chronic delivery of hypoxic gas poses both practical challenges and long-term safety concerns. We previously introduced a small-molecule, “hypoxia-in-a-pill” combining a hemoglobin affinity enhancer (GBT440) to limit oxygen delivery with a HIF-2α inhibitor (PT2399) to prevent detrimental compensatory erythropoiesis (2). Although this small-molecule regimen extended the lifespan of the *Ndufs4* KO mouse model of mitochondrial complex I deficiency and Leigh syndrome, its efficacy did not match chronic 11% FIO2. Here we optimize this regimen using GBT601, a second-generation hemoglobin affinity enhancer with longer half-life and greater hemoglobin occupancy, and show it rescues key neurodegenerative phenotypes in multiple models. When we initiate therapy in 50 day old *Ndufs4* KO mice with advanced disease, GBT601 monotherapy alleviated neurological disease phenotypes and extended median lifespan from 62 days to 105 days, while dual therapy with the GBT601/PT2399 combination extended median lifespan to 158 days. When initiated after onset of advanced disease in a mouse model of Friedreich’s ataxia, the combination halted further progression of motor phenotypes. In a mouse model of Parkinson’s disease due to a-synuclein toxicity, initiating the GBT601/PT2399 combination after onset of neurodegeneration attenuates brain hyperoxia and lipid peroxidation and reverses motor phenotypes. Importantly, the combination maintained body weight without inducing any signs of pulmonary hypertension. Our findings motivate further pre-clinical and clinical evaluation of our “hypoxia in a pill” approach for diseases with high unmet need.

## Introduction

Although seemingly counterintuitive, a mounting body of research is demonstrating that chronic moderate hypoxia can have widespread therapeutic benefits (1). Epidemiological studies have long suggested the benefits of living at high altitude. For instance, residents of Swiss (3) and Austrian (4) Alps and the western United States (5) exhibit lower mortality from cardiovascular disease, stroke, cancer, and Alzheimer’s disease. Anecdotal evidence suggests that high altitude may ameliorate motor symptoms in Parkinson’s disease (6). A notable “natural experiment” involving ∼20,000 soldiers from the Indian army stationed at high altitude (3200-4800 m, equivalent to an inspired oxygen of 11-13% at sea level) for three years revealed a lower incidence of hypertension, type 2 diabetes, and ischemic heart disease compared to that of ∼130,000 peers stationed at sea level (7). While many other environmental variables such as climate, solar radiation, diet, and activity confound these human data, preclinical research has supported reduced oxygen as a candidate driver of these benefits.

Preclinical mouse studies beginning in 2016 showed that continuous exposure to 7-11% oxygen provides profound protective effects across diverse models, including mitochondrial gray matter disease called Leigh syndrome (8–11), Leigh-like syndrome (12), optic neuropathy (13), Friedreich’s ataxia (FA) (14, 15), multiple sclerosis (16), accelerated aging phenotype (17), recovery from myocardial infarction (18) and ischemic stroke (19, 20), and Parkinson’s disease (21). The therapeutic benefits of hypoxia for rare and common forms of neurodegenerative diseases appear to be evolutionarily conserved from mice to *C. elegans* (11, 14, 21). Given the widespread role of molecular oxygen in metabolism and signaling, the therapeutic benefits of hypoxia are likely to be pleiotropic in origin, but key specific mechanisms are being identified. For example, we were the first to document that mouse models of primary or secondary mitochondrial complex I deficiency exhibit brain hyperoxia, which we have shown can be attenuated by breathing chronic, continuous hypoxia (10, 21). A low ambient oxygen also helps to preserve the activity of mitochondrial complex I (11, 14), helps to restore iron-sulfur cluster levels (14), and reduce brain lipid oxygenation (21). In most of these models, traditional antioxidants are not effective and cannot provide a simple substitute for chronic hypoxia.

Harnessing these benefits requires continuous rather than intermittent regimens of hypoxia. In the *Ndufs4* KO model of Leigh syndrome, daily 10-hour hypoxic bouts failed to provide benefit (9) while in the *shFxn* model of Friedreich’s ataxia, a 16-hour daily regimen even proved detrimental (15). We hypothesized that this failure likely stems from the elevated hematocrit and hemoglobin (Hb) induced by the hypoxic periods, which, upon return to normoxia, creates a state of excess oxygen that is detrimental. In support of this thesis, we showed that pharmacologically inhibiting HIF-2α, which blunts the erythropoietic response to low oxygen, can attenuate these detrimental effects (15).

Safely delivering chronic continuous hypoxia raises practical challenges. The sporting industry has created a variety of portable nitrogen generators, tents, and face masks, that allow athletes to train in environments simulating high altitude. We previously “repurposed” such equipment in a study of healthy volunteers to achieve normobaric hypoxia over 5 days that nadired at 11% oxygen that was safe and well-tolerated (22). It is conceivable that this equipment could be re-purposed for long-term hypoxia therapy, though this would be cumbersome.

Chronic exposure to continuous hypoxia is also associated with a number of long-term, detrimental side effects, including pulmonary hypertension (PH) and right ventricular hypertrophy (RVH) in humans (23, 24) and in mice (25). These risks are particularly pronounced during development where neonatal hypoxia interferes with lung airway and vascular development, with reduced long-term survival due to severe PH and RV failure (25, 26). Furthermore, long-term hypoxia is associated with development of paragangliomas and glomus tumors (27). Many of the detrimental side effects of chronic hypoxia are due to prolonged and sustained activation of HIF-2α (28).

These challenges underscore the need to be able to deliver hypoxia in a way that is both practical and safe, motivating us to introduce “hypoxia in a pill,” the first small molecule cocktail for inducing systemic, therapeutic hypoxia (2). Our small molecule approach was based on dual targeting of hemoglobin and HIF-2a using two classes of FDA approved drugs, respectively. First, we used the hemoglobin affinity enhancer GBT440 to decrease oxygen offloading and thereby decrease tissue oxygen levels. Linus Pauling showed in the 1940’s that oxygenated hemoglobin – corresponding to a conformation called the “R” state – has a lesser tendency to sickle (29). Small molecule hemoglobin affinity enhancers that stabilize this oxygen bound “R” conformation have been developed to treat sickle cell anemia (30, 31). However, the decrease in oxygen offloading achieved with GBT440 can in principle induce a compensatory, HIF-2α-dependent erythropoietic response which serves to increase tissue oxygen delivery and therefore limits the effectiveness of GBT440 monotherapy. Hence, we combined GBT440 with the HIF-2α inhibitor PT2399, a class of drugs that has been approved for vHL null renal tumors (2). We showed that treatment with the combination could extend the lifespan of the *Ndufs4* KO mouse model of Leigh syndrome and delay onset of neurological disease (2). A subsequent study showed that daily administration of a structural analog of GBT440 from the same initial chemical series could also extend lifespan in the *Ndufs4* KO mouse model (32), though it is an irreversible covalent binder administered at extremely high doses and never advanced into humans.

While our initial combination consisting of two FDA approved classes (2) provided proof-of-concept, it remained less effective than inhaled 11% oxygen, likely due to the short half-life and limited Hb occupancy by GBT440. Here, we sought to optimize the regimen using GBT601 (Osivelotor), a potent, second-generation, clinical-stage molecule (33–35). GBT601 exhibits greater hemoglobin occupancy and a longer half-life in rats compared with GBT440 (33). In a murine model of sickle cell disease, GBT601 treatment resulted in both an increase in hemoglobin oxygen affinity and a reduction in sickling of red blood cells (RBCs) that exceeded the effects of GBT440 preclinically. These pharmacological improvements enable equivalent HbS occupancy at a lower dose, reducing treatment burden. Importantly, GBT601 has already completed safety studies in healthy humans (34).

Here we sought to explore the therapeutic potential of GBT601 alone or the GBT601/PT2399 combination in preclinical mouse models of mitochondrial and neurodegenerative diseases in which inhalational hypoxia has previously been shown to be effective. We show that this combination is tolerated without apparent adverse effects in wild-type mice, achieves durable systemic and brain hypoxia, and when initiated in advanced stages of disease in these models, is able to alleviate neurodegenerative phenotypes. Importantly, dual targeting of hemoglobin and HIF-2a achieves these results without the detrimental cardiac side effects typically associated with chronic hypoxia.

## Results

### Dual targeting of hemoglobin and HIF-2a with GBT601/PT2399 induces systemic hypoxia

We first measured effects of daily administration of GBT601 and PT2399 on hemoglobin levels and brain tissue oxygen partial pressure (PbO2) in wild-type (WT) C57BL/6J mice. We began by treating WT mice with each drug, individually or in combination, by oral gavage daily using a fixed dose previously reported (2, 36). After three weeks of daily treatment, the average hemoglobin in vehicle-treated mice (n=5) was 14.4 +/− 0.4 g/dL, rose in mice treated with GBT601 alone to 16.5 +/− 1.0 g/dL (n=8, P=0.002) and decreased in those treated with PT2399 alone to 12.9 +/− 1.0 g/dL (n=7, P=0.02). The combination, however, led to Hb levels comparable to those with vehicle treatment (14.0 +/− 1.0 g/dL; n=8, P=0.8). These findings confirm that the erythroid response to GBT601 is blunted by the HIF-2α inhibitor PT2399 (Figure 1A).

**Figure 1.**
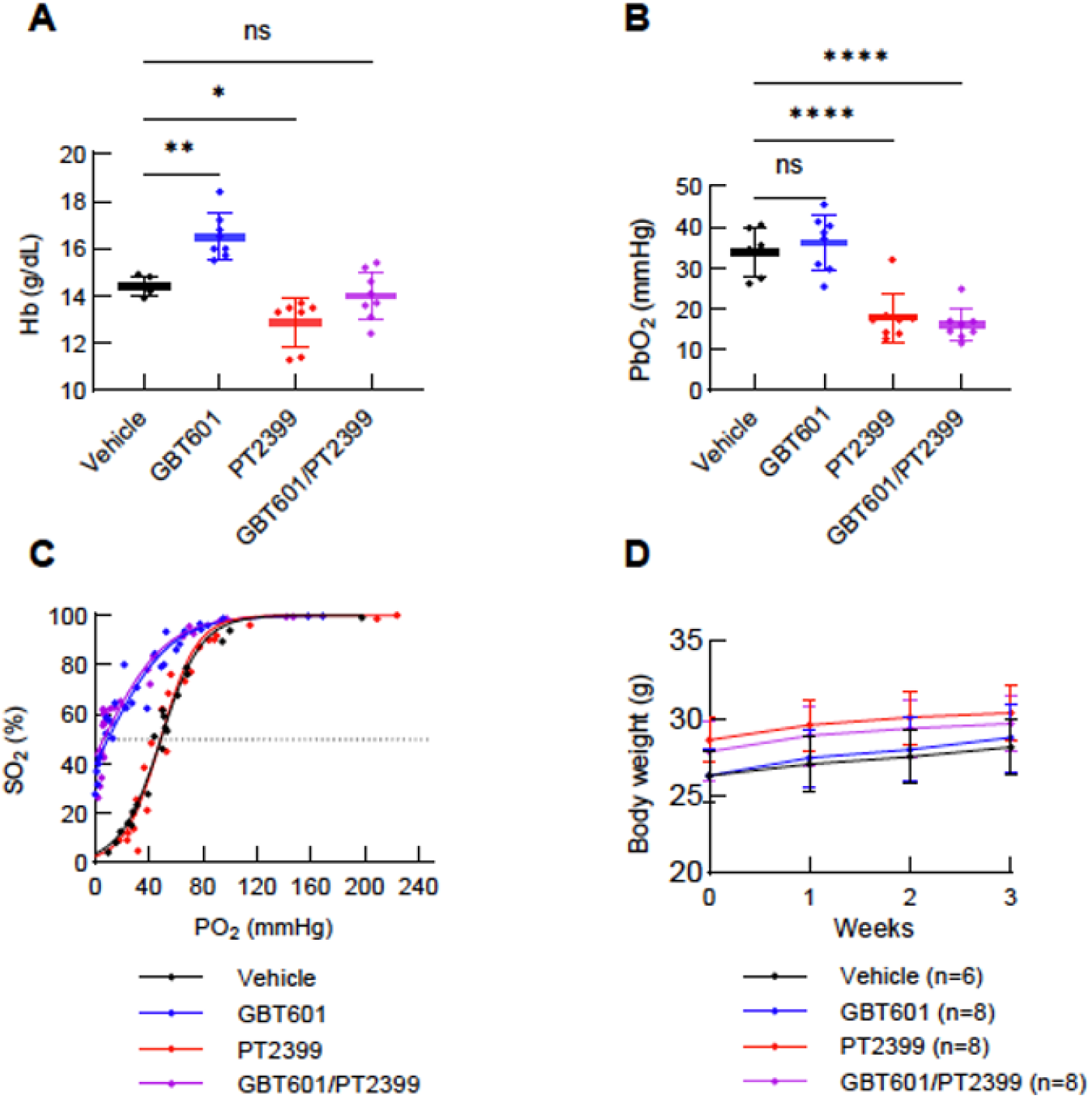
GBT601/PT2399 combination induces systemic hypoxia. (**A**) Hb and (**B**) brain PbO2 measurements in vestibular nucleus of 10-week-old C57BL/6J male mice treated with vehicle, GBT601, PT2399 or the combination for 3 weeks (n=5-8 per group). (**C**) oxygen– hemoglobin dissociation curve measurements of 12-week-old C57BL/6J male mice treated with vehicle, GBT601, PT2399 or the combination for 7 days (n=4-5 per group). (**D**) Body weight of C57BL/6J mice treated with the indicated drugs. Bar plots show the mean ± SD. *P < 0.05, **P < 0.01, ***P < 0.001, ****P < 0.0001; 1-way ANOVA with Dunnett’s test for multiple comparisons with vehicle. oxygen–hemoglobin dissociation curve compared by nonlinear regression (extra sum-of-squares F test).

To determine whether the drugs were inducing systemic and tissue hypoxia, we made brain PO_2_ (PbO_2_) measurements. Using an optical oxygen probe, we found that mice treated with GBT601 alone had similar PbO2 in the vestibular nucleus to the vehicle-treated (34.1+/− 6.1 mmHg vs. 36.2 +/− 6.8 mmHg, P=0.8), while those treated with PT2399 had significantly decreased PbO2 17.9 +/− 6.0 mmHg) (P<0.0001 vs. vehicle). The combination of GBT601 and PT2399 decreased PbO2 to 16.1 +/− 4.0 mmHg (P<0.0001 vs. vehicle) (Figure 1B).

Oxygen dissociation curves demonstrated distinct shifts among treatment groups. Vehicle-treated mice exhibited a typical sigmoidal oxyhemoglobin saturation curve with a P_50_ of 48.9 mmHg (95% CI: 47.1–50.7). PT2399 treatment alone did not substantially alter hemoglobin–oxygen affinity, with a P_50_ of 48.7 mmHg (95% CI: 45.7–51.9). In contrast, and as expected, GBT601 treatment produced a marked left shift of the oxygen dissociation curve. The P_50_ was significantly reduced to 9.6 mmHg (95% CI: 5.8–12.9), reflecting increased hemoglobin–oxygen affinity and enhanced saturation at lower PO₂ levels. The combination of PT2399 and GBT601 also demonstrated a pronounced leftward displacement, with a P_50_ of 6.4 mmHg (95% CI: 1.8–10.1), indicating preserved GBT601-mediated increases in oxygen affinity in the presence of PT2399 (Figure 1C).

Comparable to what we previously observed in mice treated with GBT440 and PT2399, the trajectory of bodyweight was the same for mice treated with vehicle, GBT601 alone, PT2399 alone and the combination of GBT601/PT2399. (Figure 1D). Thus, the combination of GBT601/PT2399 induces sustained brain tissue hypoxia and is well tolerated in wild-type mice for at least three weeks of daily treatment.

### GBT601 alone or in combination with PT2399 alleviates neurodegenerative phenotypes and extends lifespan in a mouse model of Leigh syndrome

Leigh syndrome is the most common pediatric manifestation of mitochondrial disease and is characterized by a subacute degeneration of deep gray matter regions. Patients tend to be born developmentally intact and the disease presents as a neurometabolic crisis in in infancy or in childhood with characteristic brain MRI changes. To date, more than 110 different disease genes, nearly all of which encode mitochondrial proteins, have been identified (37), and unfortunately, there are no FDA approved medicines. We previously showed that chronic, continuous exposure to 11% FIO2 initiated after weaning prevents the formation of brain pathology in the *Ndufs4* KO mouse model of Leigh syndrome due to complex I deficiency (8). As the brain disease is the cause of death in these mice, hypoxic breathing leads to a striking extension in lifespan. We later showed that when initiated after onset of advanced disease, 11% FIO2 could halt further neurodegeneration and even reverse MRI lesions and motor phenotypes (9). Moreover, we reported that the first generation “hypoxia in a pill” (2) consisting of the GBT440/PT2399 combination when initiated at the time of weaning could delay onset of disease and prolong median lifespan by 30%, whereas neither drug was effective in isolation.

Here we sought to evaluate whether the second generation GBT601 alone, or in combination with PT2399, are effective when initiated at day 50, when these mice exhibit very advanced neurological disease (Figure 2A). This is a more stringent approach (“rescue” versus “prevention”) and is more clinically relevant as most patients present after the onset of disease symptoms. Ordinarily, *Ndufs4* KO mice are born healthy, show neurological defects on day ∼35, begin to lose body weight, and succumb at ∼2 months to their neurodegenerative disease. We began therapy at day 50, when *Ndufs4* KO mice already have advanced neurological disease, and vehicle treated mice are expected to survive for only two more weeks. GBT601 alone increased median lifespan from ∼62 days (vehicle) to 105 days (P<0.0001) and maximum lifespan from 80 to 138 days (P<0.0001). Strikingly, the GBT601/PT2399 combination extended median lifespan from ∼62 to 158 days (P<0.0001) and maximum lifespan by 2.7 times from 80 to 212 days (P<0.0001, Figure 2B). Moreover, *Ndufs4* KO mice demonstrated an upward growth trajectory following initiation of treatment (Figure 2C-D). At day 70, when half of the vehicle group had died, the average body weight of the GBT601 mice was higher than the surviving vehicle treated mice (13.4 ± 1.1g vs. 10.3 ± 0.6 g, P<0.0001) (Figure 2C, SI Appendix, Fig. S1A-B). Similarly, the body weight of the GBT601/PT2399 group was higher than the mice surviving in the corresponding vehicle group (12.6 ± 1.4 g vs. 9.8 ± 0.3 g, P<0.0001) (Figure 2D, SI Appendix, Fig. S1C-D). The open field test showed that distance traveled increased from day 50 prior to treatment to day 80 with either GBT601 (P=0.01, Figure 2E) or with the combination (P=0.048, Figure 2F).

**Figure 2.**
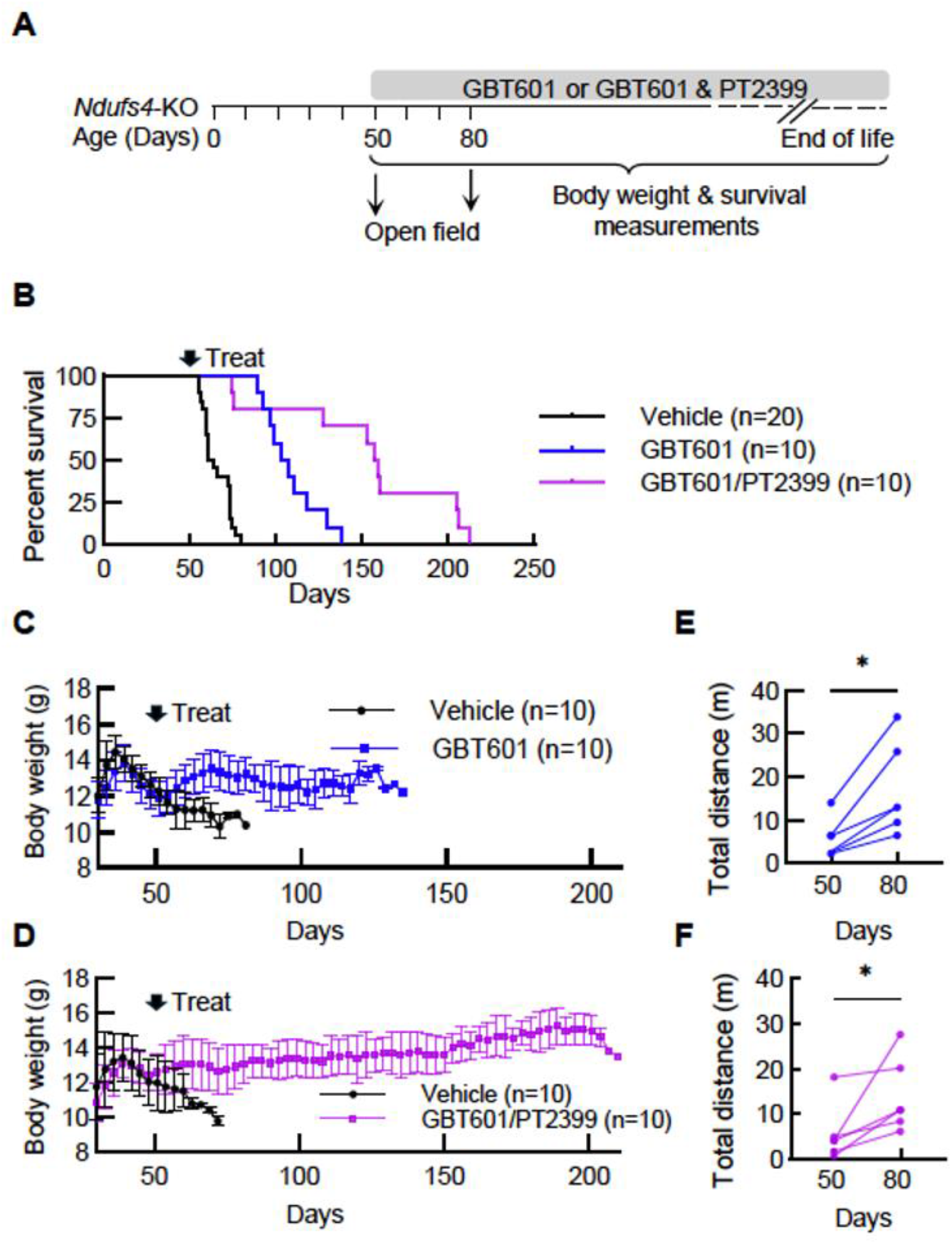
GBT601 alone or GBT601/PT2399 combination alleviates neurological disease trajectory and extends lifespan in Ndufs4 KO model of Leigh syndrome. (**A**) Schematic depicting the Ndufs4 KO mice experimental tests and time course. (**B**) Survival of Ndufs4 KO mice treated with vehicle, GBT601, or the combination. Body weight change of Ndufs4 KO mice treated with the GBT601 (**C**) or the combination (**D**). Distance traveled in 15 minutes on an open-field test of Ndufs4 KO mice treated with GBT601 (**E**) or the GBT601/PT2399 combination (**F**) (n=6 per group at E, F). Bar plots show the mean ± SD. *P < 0.05, **P < 0.01, ***P < 0.001, ****P < 0.0001; paired t test; log-rank test for survival of drug-versus vehicle–treated mice. 2 way ANOVA test for body weight of drug-versus vehicle– treated mice.

Collectively, these studies indicate that “hypoxia in a pill” monotherapy with GBT601 alone or with the dual-targeting approach GBT601/PT2399 can alleviate neurodegenerative disease phenotypes and rescue the *Ndufs4 KO* mouse when they have advanced disease, with a striking extension of survival and lifespan.

### GBT601/PT2399 halts neurological disease phenotypes in Friedreich’s ataxia mouse model

Given the striking performance of the GBT601/PT2399 combination in the *Ndufs4* KO model, we advanced it into a mouse model of Friedreich’s ataxia. Friedreich’s ataxia (FRDA) is the most common inherited ataxia due to a recessive loss of the frataxin (FXN) protein (38–40). Patients are born developmentally intact, and begin to exhibit signs of ataxia ∼10 years of age and typically develop diabetes and cardiomyopathy. Using the doxycycline-induced *shFxn* mouse model of Friedreich’s ataxia (41), we previously reported that breathing chronic, continuous 11% oxygen improved ataxia phenotypes (14). We also previously reported that initiating chronic, continuous 11% FIO_2_ for 3 weeks can rapidly reverse features of ataxia when commenced 12 weeks after doxycycline-induced *shFxn*, at which point neurological debility is advanced (15). However, in contrast to the *Ndufs4* KO mouse model (whose proximal cause of death in normoxia is the brain pathology), hypoxia did not improve lifespan in the *shFxn* model (41), where the cardiac pathology is the presumed proximal cause of death and not alleviated by hypoxia in either model (9, 14).

Here, we tested whether the GBT601/PT2399 combination could mimic the effects of inhalational hypoxia to similarly halt neurological symptoms in the *shFxn* mice once they began to exhibit signs of advanced debility (Figure 3A). As expected (41), at 24 weeks of age (after 12 weeks of doxycycline treatment), *shFxn* mice treated with doxycycline began to exhibit decreased latency to fall on rotarod testing (Figure 3B). Between 24 weeks and 27 weeks, *shFxn* mice treated with the drug vehicle alone continued to exhibit deterioration, with their latency to fall on rotarod testing declining in the surviving mice from a mean of 98 seconds to 57 seconds (paired T-test P=0.003, Figure 3C). In contrast, *shFxn* mice treated with GBT601/PT2399 better preserved their time on the rotarod, with a mean latency to fall of 90 seconds at 24 weeks and 83 seconds at 27 weeks in the surviving mice (paired T-test P=0.3, Figure 3D). At the 24 week time point, sh*Fxn* mice exhibited marked decrease in grip strength (Figure 3E) and continued to decline from a mean of 122 grams to a mean of 84 grams (paired T-test, P=0.003, Figure 3F). The GBT601/PT2399 could halt this further decline in surviving mice, with grip strength being preserved from a mean 111 grams at 24 weeks versus mean of 109 grams at 27 weeks (paired T-test, P=0.8, Figure 3G).

**Figure 3.**
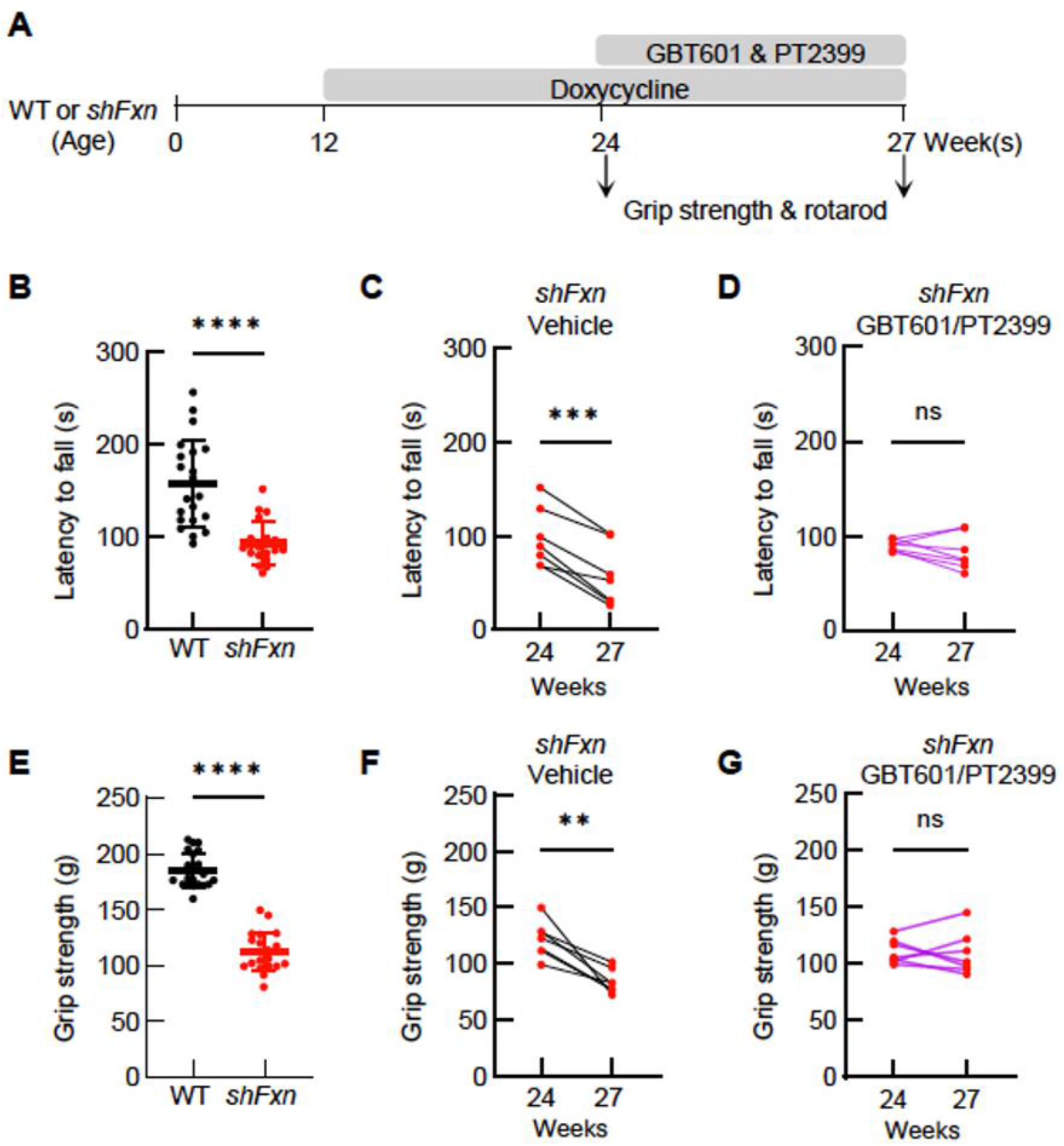
GBT601/PT2399 combination halts further neurological decline in *shFxn* mice. (**A**) Schematic depicting the *shFxn* mice experimental tests and time course. (**B**) Rotarod test and (**E**) grip strength test in 24-week-old WT and *shFxn* mice treated with doxycycline for 12 weeks (n=20 or n=21 per group at B, E). Rotarod test in *shFxn* mice treated with vehicle (**C**) or combination (**D**) at 24-week-old and 27-week-old. Grip strength test in *shFxn* mice treated with vehicle (**F**) or combination (**G**) at 24-week-old and 27-week-old (n=7 per group at C, D, F, G). Bar plots show the mean ± SD. n=group size. *P < 0.05, **P < 0.01, ***P < 0.001, ****P < 0.0001; paired t test for single comparison.

Hence, these studies indicate that initiating the combination treatment at 24 week of age, can halt further progression of the ataxia phenotype, though in contrast to 11% FIO_2_ (14), is unable to reverse these motor phenotypes.

### GBT601/PT2399 reverses motor phenotypes in the a-syn PFF-induced mouse model of Parkinson’s disease

Next we sought to determine whether our “hypoxia in a pill” combination might be effective in a mouse model of Parkinson’s disease (PD). Parkinson’s disease is the second most common neurodegenerative disease. PD is characterized by a progressive deterioration of both motor and non-motor symptoms. At a histopathological level, the hallmarks of PD are the loss of dopaminergic (DA) neurons in the substantia nigra and the formation of Lewy bodies, mainly consisting of aggregated fibrils of the α-synuclein (α-syn) protein. Secondary mitochondrial defects, notably in complex I, are frequently documented in sporadic and rare forms of this disease.

We previously reported the therapeutic benefits of breathing chronic, continuous 11% oxygen in a α-syn preformed fibrils (PFF) mouse model of PD (21). In this model, PFFs induce endogenously expressed α-syn to form aggregates closely resembling Lewy Pathology in PD, leading to significant reduction of dopamine neurons in the substantia nigra pars compacta (SNpc) (42). We showed that PFF-induced α-syn aggregation resulted in brain tissue hyperoxia, lipid peroxidation and dopaminergic neurodegeneration in the SNpc of mice breathing 21% oxygen. In mice breathing 11% oxygen, we found no difference in the abundance of α-syn aggregates, however, hypoxia attenuated the brain hyperoxia, lipid peroxidation, and DA neuron loss. Moreover, initiating hypoxia 6 weeks after PFF injection could reverse motor symptoms and halted further DA neurodegeneration (21).

We utilized a similar study design but now commenced with the GBT601/PT2399 “hypoxia in a pill” combination (instead of inhalational hypoxia) after disease onset (Figure 4A). Briefly, we administered intrastriatal injections of PFFs or monomeric α-syn (mon) as a control to 12-week-old mice to confirm onset of motor dysfunction at 18 weeks. We then treated the PFF mice or monomer mice with either GBT601/PT2399 combination regimen or vehicle from 18-week-old to 24-week-old and assessed motor behavior using the pole test (coordination, bradykinesia), the cage hang test (coordination, balance), and the open field test (gross motor and anxiety-like behavior) (43).

**Figure 4.**
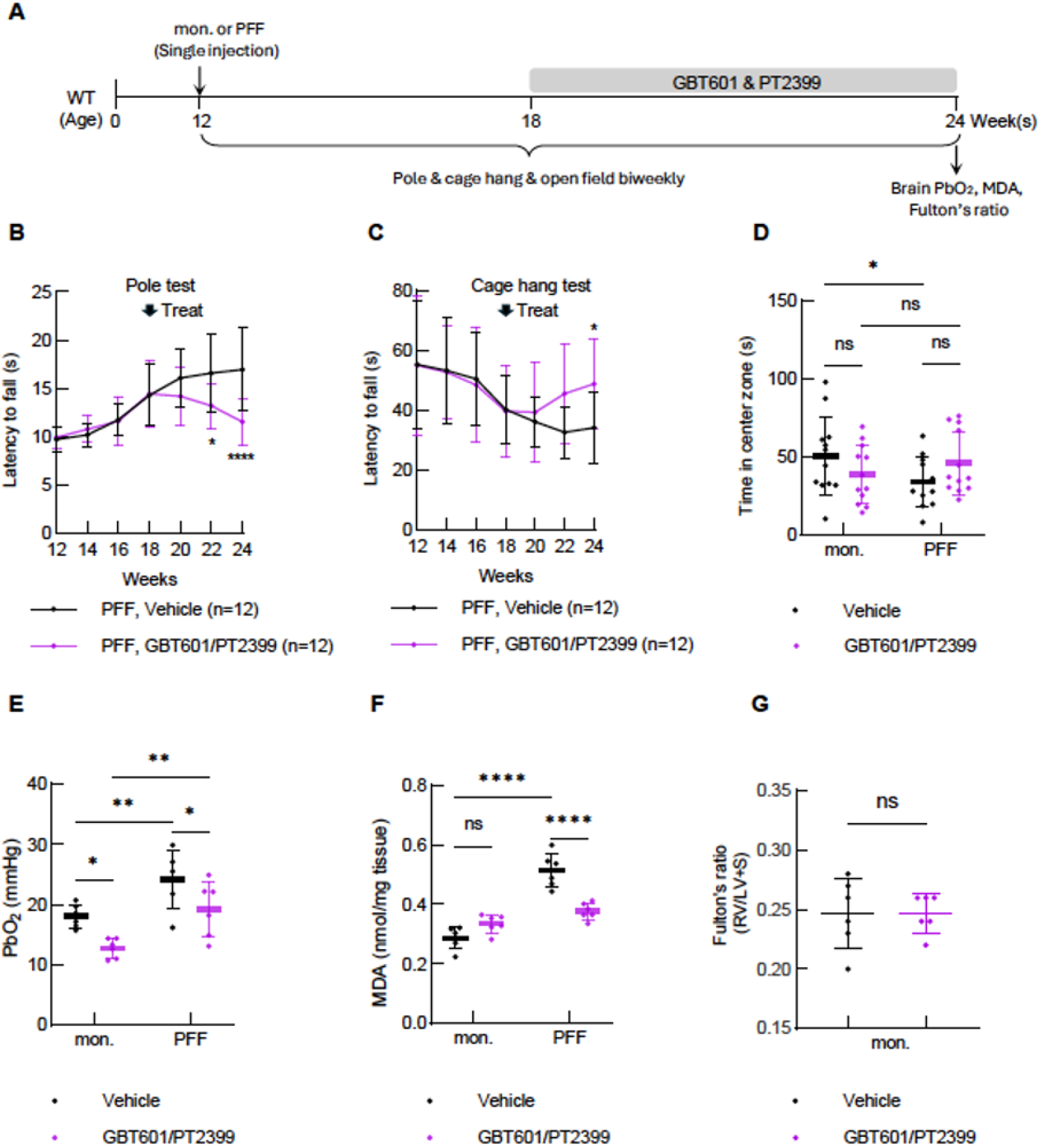
GBT601/PT2399 combination rescues α-syn-induced parkinsonism in mice. (**A**) Schematic depicting the mouse model of Parkinson’s disease experimental tests and time course. (**B**) Pole test and (**C**) Cage hang test in Parkinson’s disease (PFF) mice treated with vehicle or the GBT601/PT2399 combination. (**D**) time in center zone on an open-field test of control (mon) and Parkinson’s disease (PFF) mice treated with vehicle or the GBT601/PT2399 combination. (**E**) brain PbO2 measurements (**F**) MDA levels in SNpc tissues of control (mon) and PFF mice treated with vehicle or the GBT601/PT2399 combination. (**G**) Fulton’s ratio of control (mon) mice treated with vehicle or the GBT601/PT2399 combination. Bar plots show the mean ± SD. n=group size, n=12 per group at D. n=6 per group at E, F, G. *P < 0.05, **P < 0.01, ***P < 0.001, ****P < 0.0001. A two-way ANOVA for multiple comparisons was conducted.

Mice with injected with PFFs exhibited increased latency to descend on pole testing from a mean of 9.7 seconds at 12 weeks to 14.3 seconds at 18 weeks, suggesting development of bradykinesia. Between 18 weeks and 24 weeks, mice treated with the vehicle alone continued to exhibit deterioration, with their latency to descend on pole testing increasing from a mean of 14.3 secs (18 weeks) to 16.6 secs (22 weeks) and 16.9 secs (24 weeks). In contrast, PFF-injected mice treated with GBT601/PT2399 decreased their latency to descend on the pole test, from a mean of 14.4 secs (18 weeks) to 13.2 secs (22 weeks) (P=0.02, vs. vehicle) and 11.5 secs (24 weeks) (P<0.001, vs. vehicle, Figure 4B). Mice injected with PFFs showed marked impairment in the cage hang test. Between 12 weeks and 18 weeks, latency to fall in the cage hang test decreases from a mean of 55 seconds at 12 weeks to a mean of 40 seconds at 18 weeks. Between 18 weeks and 24 weeks, mice treated with the vehicle alone continued to exhibit deterioration, with their latency to fall on cage hang test decreasing from a mean of 40 seconds at 18 weeks to 34 seconds at 24 weeks. In contrast, PFF-injected mice treated with GBT601/PT2399 increased their mean latency to fall from 40 seconds at 18 weeks to 49 seconds at 24 weeks (P=0.03, vs. vehicle, Figure 4C). In open field testing, α-syn PFF treated mice showed decreased time in the inner zone (P=0.04) but the drug combination did not improve this phenotype (Figure 4D).

We previously reported that the α-syn PFF model exhibits brain hyperoxia and elevated levels of malondialdehyde (MDA), both of which were attenuated by chronic, continuous hypoxia breathing (21). We sought to determine whether “hypoxia in a pill” is also able to attenuate levels of brain dissolved oxygen and MDA. We measured PbO_2_ in the SNpc 12 weeks after intrastriatal injection of PFF or monomer. Mice injected with PFF exhibited higher PbO_2_ (24.1 +/− 4.7 mmHg) compared to mice injected with α-syn monomer (18.1 +/− 1.9 mmHg, P<0.01) which was attenuated by the combination treatment (19.2 +/− 4.6 mmHg, Figure 4E). Mice with α-syn aggregates exhibited higher MDA levels (0.5 +/− 0.06 mol/mg) compared to monomer-injected mice (0.3 +/− 0.04 mol/mg) (P<0.0001). The “hypoxia in a pill” drug combination attenuated this PFF-induced increase in MDA levels (0.4 +/− 0.03 mol/mg, P<0.0001 PFF vehicle vs. PFF GBT601/PT2399, Figure 4F).

Chronic hypoxia breathing is well known to drive pulmonary vascular remodeling in adults, including muscularization of small pulmonary arteries, increased pulmonary vascular resistance, and the development of pulmonary hypertension (PH) with consequent right-ventricular (RV) hypertrophy and, in severe cases, RV failure (23, 24). Fulton’s ratio—the weight of the right-ventricular free wall divided by the weight of the left ventricle plus septum—is a well-established marker of pulmonary hypertension. In mice, exposure to 11% hypoxia for 4–8 weeks typically increases this ratio from ∼0.25 to ∼0.35 (25). To determine whether the hypoxia in a pill drug combination therapy induced RV hypertrophy indicative of pulmonary hypertension (PH), we measured Fulton’s ratio in mice treated with vehicle or the GBT601/PT2399 combination for 6 weeks. Fulton’s ratio was similar in both groups (P=0.2, Figure 4G), suggesting that the drug combination did not induce PH.

## Discussion

In the current paper we have demonstrated that second generation, small molecule based “hypoxia in a pill” regimens are able to rescue disease phenotypes in multiple models of neurodegeneration. We focused on models in which we previously reported that chronic, continuous 11% FIO_2_ was effective. In the *Nudfs4* KO mouse model of Leigh syndrome, we initiated “hypoxia in a pill” therapy at day 50, when mice already have advanced disease and typically survive for less within two weeks; GBT601 monotherapy extended lifespan from 62 days to 105 days of age, while GBT601/PT2399 extended lifespan to 158 days. In the mouse model of Friedreich’s ataxia, initiating dual-targeting therapy at advanced neurological disease helps to halt further deterioration of motor phenotypes. In a model of Parkinson’s disease, initiating dual-targeting therapy after onset of advanced disease attenuates brain tissue hyperoxia and MDA levels and reverses neurodegenerative disease motor phenotypes. A compelling aspect of our “hypoxia in a pill” combination is that it appears to be tolerated without any obvious adverse effects on body weight or cardiac remodeling.

The combination appears to perform particularly well in the *Ndufs4* KO mouse model of Leigh syndrome and a-synuclein toxicity model of Parkinson’s disease, whereas the effects are more modest in the *shFxn* model of Friedreich’s ataxia. This pattern resembles what we have reported for inhalational hypoxia, too. Chronic continuous hypoxia alleviates neurological disease in the mouse models of Leigh syndrome and Friedreich’s ataxia, however, both of these models also exhibit cardiac disease, and for reasons we do not understand, cardiac disease in these models is not alleviated by hypoxia therapy. As the proximal cause of death in the *Ndufs4* KO model is their brain disease, these mice exhibit a striking lifespan extension with hypoxia therapy or as we’ve shown now, with the GBT601/PT2399 combination. In contrast, the proximal cause of death in the *shFxn* mouse model appears to be cardiac in origin and their lifespan is not extended by inhalational hypoxia or by “hypoxia in a pill” therapies. The *Ndufs4* KO mouse model and the a-synuclein toxicity model of Parkinson’s disease represent models of primary and secondary complex I deficiency, and we have documented the presence of brain hyperoxia (10, 21), which we presume is a consequence of electron transport chain deficiency. We have previously shown that this brain hyperoxia is alleviated by chronic, continuous 11% FIO_2_. Here we have again documented the presence of brain hyperoxia in the PD model, and that it is attenuated by the dual combination therapy. It may be the case that disorders associated with complex I deficiency and/or brain hyperoxia are particularly amenable to systemic hypoxia therapy. Future studies are required to determine the specific cause of death in the *Ndufs4* KO and *shFxn* mouse models treated with the “hypoxia in a pill,” and how they compare to inhalational hypoxia itself.

It is notable that our small molecule-based, dual targeting approach - rationally designed to induce tissue hypoxia while preventing its compensatory erythropoietic response - closely parallels the genetic adaptations that naturally underwent positive selection in human populations living at high altitude. The ascent to high altitude is associated with a compensatory, HIF-2α dependent erythropoiesis that strives to increase oxygen carrying capacity. This response is initially adaptive, but when prolonged can lead to detrimental side effects. Over evolutionary timescales, in certain populations such as the Tibetans, the chronic environmental pressure of high-altitude hypoxia has favored the selection of specific genetic variants in the EPAS1 (HIF-2α) pathway that blunt its activity (44–46). These adaptations prevent the maladaptive, excessive erythropoiesis typically seen in chronic mountain sickness, thereby protecting the cardiopulmonary system from the complications of hyper-viscosity. In mouse models, heterozygous loss of HIF-2α is protective against hypoxia induced pulmonary hypertension and right ventricular dysfunction (28). Our results demonstrate that this genetic mechanism that has been selected for in high altitude adapted human populations can be recapitulated pharmacologically with the HIF-2α inhibitor.

While the current work underscores the promise of a second-generation “hypoxia in a pill,” significant safety and regulatory hurdles remain. Although both drug classes are FDA approved, the first-generation HbO_2_ stabilizer GBT440 (Voxelotor) has been voluntarily withdrawn from the market due to an excess of deaths attributed to vaso-occlusive crises (VOCs) (33). This drug was initially approved on the basis of a biomarker response, and post-approval surveillance revealed continued VOCs on the drug, which is currently under investigation. We note that the physiological consequences of increased hemoglobin oxygen affinity may differ substantially between sickle cell disease -- characterized by chronic anemia, microvascular dysfunction, and abnormal hemoglobin polymerization -- and a non-sickle context with intact oxygen delivery reserve. Here, we utilized GBT601, a follow-on compound with an improved half-life and higher hemoglobin occupancy that has demonstrated a favorable safety profile in preclinical models and Phase I human trials (34, 35). Nevertheless, elucidating the etiology of class-associated VOCs is essential, and “hypoxia in a pill” regimens may ultimately be contraindicated in patients with sickle cell disease (47). Furthermore, a key caveat remains regarding the use of HIF-2α inhibitors; while FDA-approved for certain renal cancers, they frequently induce anemia, though we leverage this property to counteract the polycythemia typically induced by enhanced hemoglobin affinity. Continued pre-clinical and eventual clinical investigations of “hypoxia in a pill” regimens are vital to ensuring a responsible path toward confirming their safety, tolerability, and ultimate efficacy in diseases with high unmet medical need.

## Materials and Methods

### Animals

All animal experimentation was approved by the Massachusetts General Hospital Institutional Animal Care and Use Committee. Mice had free access to food and water and were maintained in a 12-h light–dark cycle, at a temperature between 20 and 25 °C and humidity between 40% and 60%. The design of experiments involving animals followed guidelines provided by “Animal Research: Reporting of In Vivo Experiments” guidelines.

We used 10–12-week-old, age-matched and weight-matched male C57BL/6J mice (Jackson Laboratory). To minimize variability, we used a randomized paired (matched pairs) design.

*Ndufs4* KO mice were generously provided by the Palmiter laboratory (University of Washington) (48, 49). As in our prior study (2), pups were genotyped and weaned at 21-28 d of age. Natural death or 20% body weight loss (euthanasia criteria) were used to generate survival curves. For the *Ndufs4* KO mouse experiments, data from both sexes is combined as no sex differences in survival phenotypes have been reported.

As in our prior studies (14, 15), *shFxn* mice were kindly provided by the Geschwind Laboratory (UCLA). Pups were genotyped and weaned at ∼25 d of age. Animals were balanced by age and sex and entered the study at an average of 3 months of age. Doxycycline was administered in drinking water (2 mg/mL, replaced weekly) and by intraperitoneal injection twice weekly (5 mg/kg for 10 weeks, then 10 mg/kg) (41). Body weight was monitored regularly, and mice were euthanized after a 20% loss of peak body weight.

For PD mouse studies, α-syn synthesis and purification was performed at the University of Alabama at Birmingham (50). As previously described (21), at 3 months of age, C57BL/6J mice were anesthetized with isoflurane on a stereotaxic frame (MM 8000/3, ASI instruments). Animals were unilaterally injected with 2 μl of 300 μM sonicated fibrils, or 300 μM monomeric α-syn as a control, into the right striatum. Solutions were injected at a constant rate of 0.5 μl min^−1^ using a gastight syringe (catalog #80030, Hamilton, Reno, NV) and a microinjection syringe pump (World Precision Instruments, Sarasota, FL) and the needle was left in place for 2 min; this was followed by slowly withdrawing the needle. The coordinates for the right striatum were +0.2 mm anteroposterior (AP), ±2.0 mm mediolateral (ML) and −3.4 mm dorsoventral (DV). All mice survived after stereotaxic injection of α-syn monomer or PFF and were used for the following experiment.

### Drug Administration

We purchased GBT601 and PT2399 from MedChemExpress and used dosing regimens as previously described (2, 36). Mice were treated via oral gavage daily at a dose of 100 mg/kg of PT2399 and 150 mg/kg of GBT601. Vehicle-treatment refers to either the vehicle for PT2399 (10% ethanol, 0.5% methyl cellulose/phosphate buffered saline) or the vehicle for GBT601 (10% DMSO, 0.5% methyl cellulose/phosphate buffered saline).

### Hemoglobin measurements

As we reported previously (2, 15), blood samples (80 μL) were obtained by tail snip and collected into heparinized capillary tubes. Hemoglobin concentration were measured using an ABL800 FLEX blood gas analyzer (Radiometer, Copenhagen, Denmark).

### Brain tissue PO_2_ measurement

Brain tissue PO_2_ was measured as previously (2, 21). For PbO_2_ measurements in the vestibular nucleus, mice were anesthetized using isoflurane (induction at 2–4%, maintenance at 1.5%), intubated, and ventilated mechanically using a tidal volume of 8 ml/kg, a respiratory rate of 110 breaths per minute, and FIO_2_ of 21%. For PbO_2_ measurements in the SNpc, mice were anesthetized with isoflurane (induction at 2–4%, maintenance at 1.5%) with FIO_2_ of 21% via a nose cone. Mice were placed in a prone position and the head stabilized using a stereotaxic frame (ASI Instruments, MI). Following skin incision and dissection, the skull was opened using a micro-drill (MD-1200, Braintree Scientific, MA) and a PO_2_ probe was inserted at the desired location. The coordinates were ML = −1.25 mm, AP = −6.00 mm, and DV = −3.90 mm or ML = 1.20 mm, AP = −3.20 mm and DV = −4.40 from the bregma for vestibular nucleus or SNpc, respectively. We measured brain tissue PO_2_ using an optical PO_2_ probe (OxyLab, Oxford Optronix, Abingdon, UK). During these brain PO_2_ measurements, we reduced the depth of anesthesia by decreasing Isoflurane concentration to 1% to minimize the effects of anesthesia.

### Oxygen Dissociation Curve Determination

Twelve-week-old mice received daily oral gavage with vehicle, PT2399, GBT601, or the combination of PT2399 and GBT601 for 7 consecutive days (n = 4-5 mice per group). Two to three hours after the last administration, mice were anesthetized with isoflurane, underwent tracheostomy, and were mechanically ventilated. The right carotid artery and right jugular vein were cannulated with PE-10 catheters for arterial and venous blood sampling, respectively. To generate oxygen dissociation curves, inspired oxygen fraction (FiO₂) was stepwise adjusted to achieve a broad range of arterial and venous oxygen tensions (PaO₂). At each FiO₂ level, arterial and/or venous blood gases were obtained. Blood samples were analyzed immediately using a blood gas analyzer (ABL90 FLEX PLUS Radiometer). Oxygen saturation values used for curve construction were calculated as functional hemoglobin oxygen saturation (SO₂) to account only for hemoglobin species capable of binding oxygen. Functional saturation was calculated as:

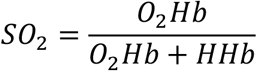

Where HHb represents deoxyhemoglobin and is calculated as: 100 – O2Hb – COHb – MetHb. Oxygen dissociation curves were constructed by plotting functional SO₂ against corresponding PaO₂ values. Data were analyzed using GraphPad Prism. Curves were fitted by nonlinear regression using a four-parameter logistic equation (log[agonist] vs. response). The upper and lower plateaus were constrained to 100% and 0% saturation, respectively, while P50 and Hill slope were allowed to vary freely.

### Accelerating rotarod measurements

We performed rotarod measurements as previously reported (14, 15), using a device available from Ugo Basile. Mice were acclimated to the testing room for 30 min prior to assessment. The rod accelerated at 5 rpm/min to a maximum speed of 40 rpm. Each testing session consisted of three trials separated by a minimum 10-min rest period, and the median latency to fall was recorded. If mice used their body to grasp the rod (rather than walking on it) for more than 10 sec, this time was recorded as time of fall.

### Grip strength test

Forelimb and hindlimb grip strength were measured using a grip strength meter (Bioseb). Each mouse completed three consecutive efforts with 30 seconds of rest in between measurements. The highest force generated (g) was recorded. If the time period of gripping was less than 10 sec, that experiment was not included. Each mouse was tested three times, and the trials were averaged.

### Open field test

We performed open field testing as previously reported (2), The open field apparatus consisted of four open-field boxes measuring 50×50 cm. Each box was surrounded by opaque black walls, and the whole apparatus was shielded from outside influences by walls with a height of 1.5 meters. After an acclimation period of 60 minutes inside the procedure room, one mouse was placed in the center of an openfield box. Up to four mice were tested in the apparatus at once. The animals were allowed to explore the field freely for 15 minutes while being recorded by the ANYaze software with an overhead camera (v7.3, Stoelting Co., Wood Dale, IL). The position and movement of the animals were automatically tracked and analyzed by the ANY-maze software. The experimenters were blind to the treatment groups and, after starting the recording, left the room for the duration of the test. Before and after each test run, 70% ethanol was used to clean the apparatus.

### Pole test and cage hang test

We performed behavioral tests as previously reported (21). Behavioral tests were conducted between 8:00 and 12:00 for the pole test and the cage hang test or 12:00 and 17:00 for the open field test. To conduct the pole test, a vertical metal pole with 2-cm diameter and 40-cm length was placed on the floor of a housing cage. Mice were placed on the top of the vertical metal pole and the time needed to reach the floor was measured. To conduct the cage hang test, a metal wire frame of the housing cage (WBL7115SMD-AMG, Allentown) was placed approximately 40 cm above the cage floor. Mice were hung on the metal wire frame and the time needed to drop onto the cage floor was measured. Each test was conducted five times and the average of the last three trials was used for the analysis.

### Detection of MDA

We measured the concentration of MDA in the SN tissue to evaluate the effect of drugs using a commercially available kit (cat. no. ab118970, Abcam). The assay was conducted according to the manufacturer’s protocol.

### Fulton’s Ratio Measurement

After euthanasia, mouse hearts were excised from the thoracic cavities, the atria removed, and the free wall of the right ventricle was separated from the left ventricle and the septum. The ratio of the weight of the right ventricle to the weight of the left ventricle and the septum was calculated (Fulton’s ratio). All measurements were performed by an investigator blinded to the treatment groups.

### Quantification and statistical analysis

Data are reported as mean ± SD. Statistical analyses were performed using GraphPad Prism 10.6.1 software. Two-way ANOVA test was used for multiple comparisons. 1-way ANOVA with Dunnett’s test for multiple comparisons with vehicle. Paired T-test for single comparisons for drugs to vehicle. P-value < 0.05 was considered to indicate statistical significance. A log-rank test for survival of drug-versus vehicle–treated mice. oxygen–hemoglobin dissociation curve compared by nonlinear regression (extra sum-of-squares F test).

## Acknowledgments

We thank Rob Rogers and Maria Miranda for feedback on this project and the manuscript. This work was supported by the NIH (R01NS112373 to FI and R01NS124679 to VKM); the Marriott Family Foundation; the Abby Mac Foundation; the Daniel Garland Fund; Friedreich’s Ataxia Research Alliance (to VKM). LZ was supported by the NIH T32 Research Training Grant (T32-GM007592) and the Foundation for Anesthesia Education and Research Mentored Research Training Grant (FAER MRTG). VKM is an Investigator of the Howard Hughes Medical Institute.

