## Supplemental Figure 1 for "A second-generation “hypoxia in a pill” rescues neurodegenerative phenotypes across distinct mouse models"

Fig. S1.

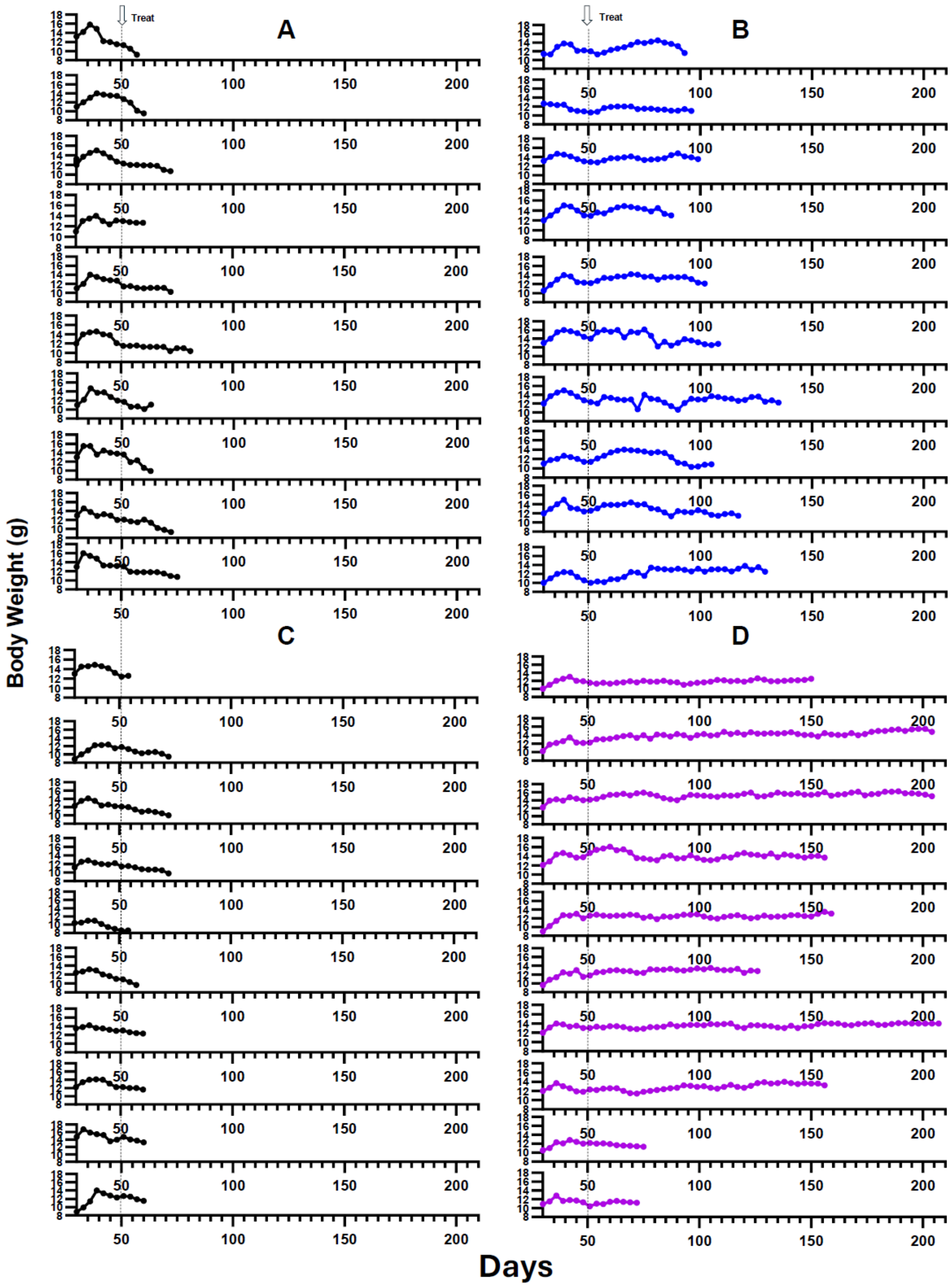


**Fig. S1. Effects of "hypoxia in a pill" regimens on growth trajectory of *Ndufs4* KO mice.**

Individual mouse body weight trajectories for *Ndufs4* KO mice treated with (**A**) vehicle, (**B**) GBT601, (**C**) vehicle, or (**D**) GBT601/PT2399.
